# BoCluSt: bootstrap clustering stability algorithm for community detection in networks

**DOI:** 10.1101/008656

**Authors:** Carlos Garcia

## Abstract

**Background:** The identification of modules or communities of related variables is a key step in the analysis and modelling of biological systems. Many module identification procedures are available, but few of these can determine the module partitions best fitting a given dataset in the absence of previous information, in an unsupervised way, and when the links between variables have different weights. Here I propose such a procedure, which uses the stability under bootstrap resampling of different alternative module structures as a criterion to identify the structure best fitting to a set of variables. In its present implementation, the procedure uses linear correlations as link weights.

**Results:** Computer simulations show that the procedure is useful for problems involving moderate numbers of variables, such as those commonly found in gene regulation cascades or metabolic pathways, and also that it can detect hierarchical network structures, in which modules are composed of smaller sub modules. The procedure becomes less practical as the number of variables increases, due to increases in processing time.

**Conclusions:** The proposed procedure may be a valuable and robust network analysis tool. Because it is based on comparing the amount of evidence for different module partitions structures, this procedure may detect the existence of hierarchical network structures.

## Background

Complex systems are often modelled and analysed as networks of related elements (nodes) connected by edges representing the relationship between them [1]. In Biology, many of these networks show a modular structure: nodes can be grouped into communities or modules so that there is a dense web of edges among nodes in the same module and a thin one between nodes in different modules [2–5]. Such structure, or modularity, has been observed in gene expression networks [6, 7], protein-protein interactions [8], metabolic [9] [10] and developmental [11, 12] pathways, and species interactions in ecosystems [13, 14].

The building of models for the study of network structure, function, regulation or evolution may require the use of module identification (also called community detection) procedures. Many such procedures have been proposed. Some are confirmatory, requiring a prior knowledge of module demarcations or at least of the number of modules present (e.g., [15, 16]). Other procedures (e.g., [17–23]) are exploratory and unsupervised, making no assumptions about modules. They use a previously known set of edges between nodes to identify the partitions among nodes that maximize some criterion of modular structure. Typically, they do not consider variation in the strength of the links between different nodes, i. e., they are unweighted. This may of course entail a loss of relevant information, as the heterogeneity in edge weights may be fundamental for the understanding of the whole network [24]. There are finally exploratory, unsupervised procedures considering different weights for each edge. In this category are the procedures of Rosvall and Bergstrom [25], based on simulated annealing and Blondel et al. [26] based on the maximization of modularity, defined as the number of edges falling within modules minus the number expected if edges were placed at random [19]. These two procedures are fast and applicable to very large networks [27], but they do not take into account the degree of precision in the estimation of these edges’ weights. This may be not critical when the focus is on the identification of large-scale patterns in big data sets, but might become an important limitation in smaller problems as those typically found in the analysis of gene regulation pathways, where the basic aim is to determine the location of variables into particular modules. In these situations it can be important to consider the robustness of module allocations, which would depend heavily on the precision in the estimation of edges weights [24].

Here I propose a new procedure combining a clustering algorithm with bootstrap resampling to identify modules of correlated variables measured in the same individuals. In cluster analysis terminology, these modules would be R-mode (because it is the variables, not the measured individuals that are being grouped [28]) variational (because the edges consist on correlations between the variables, which are represented as nodes [29]) clusters. The procedure takes into account that the correlations constituting the network edges may vary and their value may have been estimated with limited precision. I use computer simulation to show that it is superior to the procedure of Blondel et al. in the identification of variational modules in data sets with a moderate number of variables, and also that it can detect the existence of hierarchical module structures.

## Implementation

For an *n*-variables dataset, a clustering method (in the present implementation of the procedure, k-means clustering, based on the R *kmeans* function) is applied to obtain partitions into 2 to *n-1* clusters. A vector of variables’ coincidences **c** of length *(n*^2^*-n)/2* (i.e., the number of non-redundant pair wise combinations of variables) is obtained for each of these *n-2* cluster analyses, with values of 1 if the two corresponding variables were assigned to the same cluster in that analysis and of 0 otherwise. Now the stability of each of the *n-2* analyses is tested by bootstrap resampling of the individuals’ observations in the original dataset. For each resample, the above *n-2* cluster analyses are done and the corresponding **c** vectors obtained. These vectors are then compared across bootstrap samples.

If a real, detectable module structure existed in the data, bootstrap-replicated cluster analyses considering the real number of modules-clusters would tend to allocate variables in the same clusters, so that the variance across resamples would be low for each element in **c**.

A given pair of variables would tend to be either in the same cluster, the corresponding **c** value being one in most of resamples, or in different clusters, the **c** tending to be zero. In analyses considering wrong numbers of clusters -or analyses of data with no community structure-, each bootstrap replicate would result in clusters containing random combinations of variables, and the **c** values variance across bootstraps would be higher. In the procedure proposed here, the variances between resamples are calculated for each element of **c** and number of clusters, and the number of cluster partitions with the minimum value for the sum of these *(n*^2^*-n)/2* variances (i.e., that resulting in the most stable **c** vectors) is selected as the best estimate of community structure in the original data. Figure 1 illustrates the basic framework for this approach. The sum of variances can be used to compare the results obtained for the different numbers of clusters.

**Figure 1.**
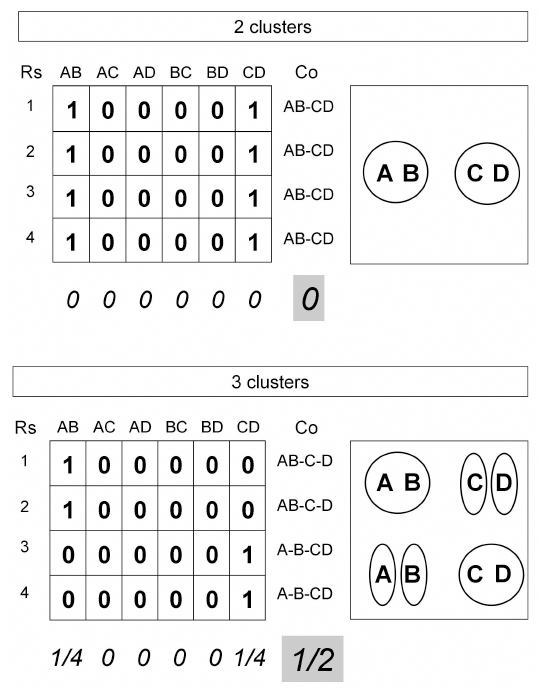
Community structure detection in a four variables (A to D) example. Cluster analyses considering two (up) and three clusters (down) are applied to four resamples (Rs) drawn from the same original data set. In each resample, each pair of variables is given a value of one if the two variables were allocated to the same cluster and zero otherwise. When the number of clusters considered in the analysis corresponds to the true community structure in the data (the two clusters case in this example), variables tend to be allocated to the same clusters in different resamples (Co – cluster composition). Thus, the above 0/1 values recording the presence/absence of coincidences have low variances (in cursive) when calculated within pairs of variables (columns in the arrays). Variances increase when other cluster numbers are considered (three in this example) because allocations become less stable under resampling. The sum of **c** variances (on a grey background) will be minimum for the number of clusters best fitting the true community structure.

It must be taken into account however that the distribution of this sum of variances is not independent of the number and size of clusters considered in the successive *n-2* analyses. To correct for this effect, the sums are made relative to their expected values in a null situation with the same number of clusters and a lack of correlation between variables. The result is the variance criterion used below. The null situation values are obtained by randomizing the observed variable values independently across individuals. Thus, while the univariate distributions are maintained, any correlation between variables disappears.

I studied the performance of the proposed method in simulated datasets of grouped variables *x*_*ij*_:

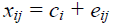

where *c_i_* was common to all *x* variables in module *i* and caused correlation among them, and *e*_*ij*_ was specific to each *x*. The considered datasets differed in number of variables, distribution of module sizes, total number of observations, correlation between variables in the same module and variables distributions (Table 1).

**Table 1.** Cases considered in Figure 2. These differed in number and distribution of variables, number and sizes of modules (there were as many modules as sizes listed), sample sizes, variables distributions and variance of the components *c* common to variables in the same correlated group. In all cases, every variable included an independent component *e* with variance = 0.050. The correlations corresponding to the three considered variances of *c* were 0.375, 0.231 and 0.167. * *c* was generated as a beta variable with parameters α = 0.246 and β = 2, and *e* (see main text) as a beta variable with α = 0.625 and β = 2,using R function *rbeta;* the resulting *x, c* and *e* distributions were markedly asymmetric. ** *c* was generated as a uniform variable with range 0 to 0.600 and *e* as a uniform variable with range 0 to 0.775 using R function *runif*

| Case | Variables<br>number | Modules<br>sizes | Sample size | Components<br>distribution | $v(c)$ |
| --- | --- | --- | --- | --- | --- |
| a | 8 | 4, 4 | 100 | Normal | 0.030 |
| b | 8 | 2, 2, 2, 2 | 100 | Normal | 0.030 |
| c | 8 | 4, 4 | 100 | Normal | 0.010 |
| d | 8 | 4, 4 | 100 | Normal | 0.015 |
| e | 8 | 4, 4 | 25 | Normal | 0.030 |
| f | 8 | 4, 4 | 50 | Normal | 0.030 |
| g | 8 | 1,1,1,1,1,1,2 | 100 | Normal | 0.030 |
| h | 8 | 5, 2, 1 | 100 | Normal | 0.030 |
| i | 8 | 4, 4 | 100 | Beta | 0.030* |
| j | 8 | 4, 4 | 100 | Uniform | 0.030** |
| k | 4 | 2, 2 | 100 | Normal | 0.030 |
| l | 16 | 8, 8 | 100 | Normal | 0.030 |

I studied the ability of the proposed method to detect hierarchical correlation structures (i.e., the presence of sub-modules within modules) by simulating datasets with variables:

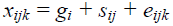

where *g*_*i*_, *S*_*ij*_ and *e*_*ijk*_ are module, sub-module and variable-specific effects.

## Results and discussion

In the non-hierarchical cases, the proposed procedure was able to identify the correct number of modules even for small size samples and moderate correlations between variables in the same module (Fig. 2). Thus, a sample size of 25 (Fig. 2e) was enough to easily identify two modules of four variables having a correlation of 0.375, and modules of variables having a correlation of 0.231 were easily detected using samples of size 100 (Fig. 2d). The performance of the procedure did not obviously depend on module number and size, the homogeneity of these sizes (Fig. 2g, 2h) or the variables’ distributions (Fig. 2i, 2j). The less favourable situations were those with the lowest correlation within modules (0.167, Fig. 2c) and the lowest number of variables (four variables, Fig. 2k). In the latter case, the variance criterion was clearly under the corresponding value for the null case, but the differences between the two and the three clusters solutions were very slight.

**Figure 2.**
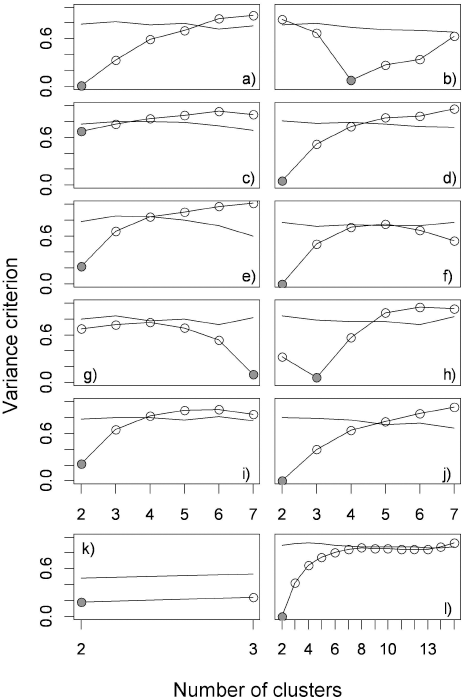
Values for the variance criterion (circles) in the cases listed in Table 1 (a single, randomly taken simulated data set per case, with 100 randomized null data sets and 500 bootstrap resamples per data set), along with the lower 2.5 percentile (simple lines) for the corresponding null situation of no correlation between variables. Grey circles mark the value for the true number of modules in the original sample.

The proposed procedure detected was able to detect hierarchical modular structures, especially when the hierarchy was regular, i.e., the pattern of subdivision was the same in all clusters (Fig. 3a, 3d and 3e). These regular partitions appeared as local minima in the sum of variances profile: two and four clusters in Figure 3a; two, four and eight clusters in Figure 3d. The procedure failed in the case of four modules and eight sub-modules (Fig. 3e) in which there the second local minimum was found for nine clusters, instead of eight. This suggests that correct community identification might require larger sample sizes as datasets become less structured and the number of independent modules increases.

**Figure 3.**
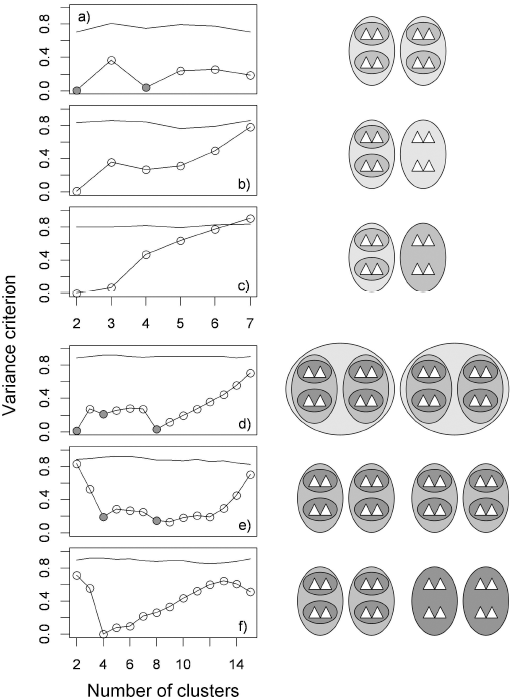
Analysis of hierarchical communities (a single, randomly taken simulated data set per case, with 100 randomized null data sets and 500 bootstrap resamples per data set). The graphs (left) show the variance criterion for all possible numbers of clusters along with the lower 2.5 percentile for the corresponding null situation of no correlation between variables (simple lines). Grey circles mark correct clustering results for regular partitions. The diagrams to the right represent the different situations. The grayscale indicate the value of the correlation between variables (triangles) in the same ellipse. These were 0.273 and 0.545 in the eight variables cases (a − c), and 0.214, 0.429 and 0.643 in the 16 variables cases (d to f).

Defining a single correct result becomes harder for less regular partitions. For example, in Figure 3b two or three clusters could be identified. While the partition into two modules was easily identified, that into three modules resulted in a local maximum instead of a minimum. This maximum disappeared when the correlation between variables in the large module in the right of the diagram increased (Fig. 3c), which, not unexpectedly, suggests that community detection is easier when edges within these communities are strong. In any case, the low criterion values for two and three clusters seen in Figure 3c would not be unambiguous evidence of hierarchical clustering, because the criterion values neighbouring a minimum can be also low in non-hierarchical situations (see for example Fig. 1h and Fig. 1k).

Figures 3e and 3f show many consecutive low values for the variance criterion. This could be in relation with the fact that many partitions are possible in these cases. For example, partitions into four, five, six or eight clusters could be possible in Figure 3e.

However, this could not explain all results. The criterion values remained low beyond eight, the last “correct” number of clusters. In any case, a comparison of Figures 2 and 3 suggests that profiles showing several points of inflexion could be indicators of hierarchical modular structures.

I made multi-sample simulations to compare the proposed procedure with that of Blondel et al. (for the latter I used the R CRAN package *igraph* [29]). Neither procedure ever failed to identify two modules for sample sizes of one hundred and moderate correlations of 0. 375 (Fig. 4 2C3). The Blondel et al. procedure was somewhat better than that proposed here when the correlation was reduced to 0.167 (Fig. 4 2C1). However, it was clearly worse in the case of four modules, as it failed to find four clusters as the most frequent result when the correlation was 0.375 (Fig. 4 4C3) and completely failed to detect them when the correlation was 0.167 (Fig. 4 4C1). In the same situations, the right solution of four clusters was that most frequently found by the proposed procedure.

**Figure 4.**
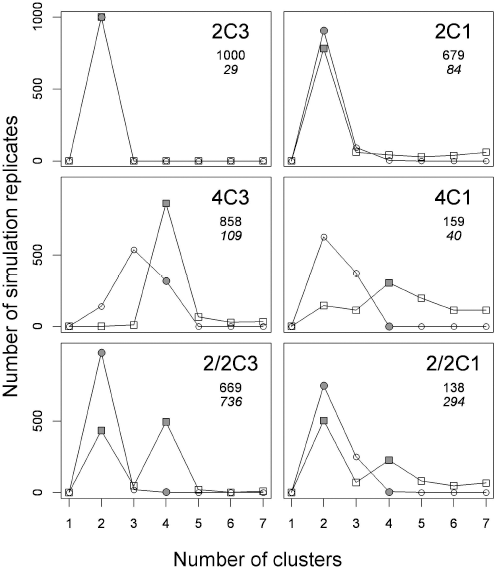
Frequency of numbers of clusters detected in computer simulations (1000 replicates, eight variables). 2C and 4C are cases with two and four variable modules respectively, and 2/2C, hierarchical situations of two modules each divided in two sub­modules. The numbers to the right of the Cs mark the variance of components *c,* common to variables in the same module or sub-module: 1, variance = 0.01; 3, variance = 0.03 (the variance of component *e* was = 0.05 in all cases). Circles and squares show results for the proposed and the Blondel et al. procedures respectively. Grey symbols mark the value for the true number(s) of clusters. Results for one cluster are given in the x axe to ease the graphs’ interpretation. This could never be the number of clusters detected because the results for the simulated and randomized cases are always identical in that case: all variables in the same and only cluster. The smaller font values are the numbers of replicates finding correct and significant results. In the hierarchical cases, this is two significant minima in the variance criterion for two and four clusters with the proposed procedure. In italics, number of replicates finding the same two minima, whether significant or not.

In the hierarchical cases, the Blondel et al. procedure found only two clusters in an overwhelming majority of replicates, whereas the proposed procedure found two and four clusters as the most frequent solutions. For individual replicates, the proposed procedure would detect hierarchical situations as multiple minima for the variance criterion, as in Figure 3a. The proposed procedure was able to detect the hierarchical structure in most replicates when the correlation was moderate (Fig. 4 2/2C3, in italics), but only in a minority when the correlation was low (Fig. 4 2/2C1).

The proposed procedure was better than the Blondel et al. procedure in most simulations made here, especially for the cases involving the smallest module sizes. The low performance of the Blondel et al. procedure was likely related to the modularity-based methods’ “resolution limit in community detection”. This limit is most likely to occur when the number of module internal links is of the order of the square root of twice the total number of links in the network or smaller (Fortunato and Barthelemy 2007), i.e., when modules are small. The proposed procedure seems to be unaffected by that limit, since it can easily detect two-variables modules (see Fig. 1b). However, the use of bootstrap resampling by this procedure might command too many computational resources to be practical for the analysis of large sets of variables, as those in genome-wide or human social networks. Thus, the proposed procedure could be a useful complement for low computational complexity, large-scale procedures such as that of Blondel’s et al., especially when small modules are involved. The two procedures would not be equivalent for any problem size, however, because they do not use the same kind of information. Instead of starting with a previously known set of edge weights, the procedure proposed here simultaneously estimates both the weights and the community structure. The approach of measuring the consistency of a found community structure was already proposed by Duch and Arenas [21]. They used an extremal optimization algorithm that could result in different network partitions in different runs, so that they could calculate the fraction of times a pair of nodes was allocated to the same module. However, they did not use consistency as a criterion to identify the optimum community structure among a set of possible structures, as done here.

Figure 2 considers only from 2 to n-1 as possible cluster numbers. This is because considering coincidences in module allocation (and it could be argued that the very idea of clustering) does not make sense when there are n clusters of size one (and therefore no coincidences) or there is a single cluster including all variables (total coincidence). However, because the proposed procedure compares the obtained results with those expected under the absence of community structure, it is possible to detect this absence: if none of the 2 to n-1 partitions were below the lower 2.5 percentile of the null distribution. In such a case, it would be concluded that there is no community structure. Thus, the proposed procedure does not only provide an estimate of the number of modules, but also of the reliability of that estimate and of the overall degree of structure in the data. It also makes it possible to compare the reliability of alternative solutions.

It must be noted that, while being able to detect some hierarchical modular structures, the proposed procedure does not provide a formal diagnostic for such structures. As seen in the results section, some of the variance criterion results could correspond both to hierarchical and no-hierarchical structures. These structures are more easily detected when the hierarchy is regular, in the sense that variables groups are composed of the same number of subgroups, of the same size and the same correlation between variables. This tends to result in separate minima for the variance criterion, which is characteristic of hierarchical structures.

The present formulation of the proposed procedure uses correlations as distance measures between variables and K-means as the clustering algorithm, but its approach of evaluating alternative partitionings based on measuring its consistency in the face of resampling would be compatible in principle with any combination of distance definitions and clustering algorithms.

## Conclusions

The proposed procedure could be a useful tool for the analysis of networks of small to moderate size, making it possible to get an unsupervised estimate of the number of clusters present.

## Availability and requirements

BoCluSt is available as an R function in Sourceforge: http://sourceforge.net/projects/boclust/files/BoCluSt.txt/download

## Competing interests

I declare no competing interests

## References

1. Proulx SR, Promislow DE, Phillips PC: Network thinking in ecology and evolution. Trends Ecol Evol 2005, 20: 345–353.

2. von Dassow G, Munro E: Modularity in animal development and evolution: elements of a conceptual framework for Evo-Devo. J Exp Zool B Mol Dev Evol. 1999, 285: 307–325.

3. Callebaut W, Rasskin-Gutman D. Modularity: Understanding the Development and Evolution of Natural Complex Systems. Cambridge: MIT Press; 2005. 455p.

4. Wagner GP, Pavlicev M, Cheverud JM: The road to modularity. Nat Rev Genet, 2007, 8:921–931.

5. Hintze A, Adami C: Evolution of complex modular biological networks. PLoS Comput Biol 2008, 4(2): e23

6. Segal E, Friedman N, Koller D, Regev A: A module map showing conditional activity of expression modules in cancer Nat Genet 2004, 36: 1090–1098.

7. Litvin O, Causton HC, Chen BJ, Pe’er D: Modularity and interactions in the genetics of gene expression. P Natl Acad Sci USA 2009, 106: 6441–6446.

8. Spirin V, Mirny LA: Protein complexes and functional modules in molecular networks. P Natl Acad Sci USA 2003, 100: 12123–12128.

9. Ravasz E., Somera AL, Mongru DA, Oltvai ZN, Barabási A: Hierarchical organization of modularity in metabolic networks. Science 2002, 297:1551–1555.

10. Kreimer A, Borenstein E, Gophna U, Ruppin E: The evolution of modularity in bacterial metabolic networks. P Natl Acad Sci USA 2008, 105: 6976–6981.

11. Wagner GP: Homologues, natural kinds, and the evolution of modularity. Am. Zool 1996, 36: 36–43.

12. Bolker JA: Modularity in development and why It matters to Evo-Devo. Amer. Zool 2000, 40: 770–776.

13. Jordano P: Patterns of mutualistic interactions in pollination and seed dispersal: connectance, dependence asymmetries, and coevolution. Am Nat 1987, 129: 675–677.

14. Dupont YL, Olesen JM: Ecological modules and roles of species in heathland plant-insect flower visitor networks. J Anim Ecol 2009, 78: 346–353.

15. Klingenberg CP: Morphometric integration and modularity in configurations of landmarks: tools for evaluating a priori hypotheses. Evol Dev 2009, 11:405–421.

16. Fruciano C, Franchini P, Meyer A: Resampling-Based Approaches to Study Variation in Morphological Modularity. PLoS ONE 2013, 8:e69376.

17. Girvan M, Newman MEJ: 2002. Community structure in social and biological networks. P Natl Acad Sci USA 2002, 99: 7821–7826.

18. Clauset A, Newman MEJ, Moore C: Finding community structure in very large networks. Phys Rev E 2004, 70:066111.

19. Newman MEJ: Modularity and community structure in networks. P Natl Acad Sci USA 2006, 103: 8577–8582.

20. Guimera R., Nunes Amaral LA: Functional cartography of complex metabolic networks. Nature 2005, 433:895–900.

21. Duch J, Arenas A: Community detection in complex networks using extremal optimization. Phys Rev E 2005, 72:027104.

22. Sales-Pardo M., Guimerã R, Moreira AA, Amaral LAN: Extracting the hierarchical organization of complex systems. P Natl Acad Sci USA 2007, 104: 15224–15229.

23. Reichardt J, Bornholdt S: Partitioning and modularity of graphs with arbitrary degree distribution. Phys Rev E 2007, 76: 015102.

24. Barrat A, Barthélemy M, Pastor-Satorras R, Vespignani A The architecture of complex weighted networks. P Natl Acad Sci USA 2004, 101:3747–3752.

25. Rosvall M, and Bergstrom CT Maps of random walks on complex networks reveal community structure. P Natl Acad Sci USA 2008, 105:1118–1123.

26. Blondel VD, Guillaume JL, Lambiotte R, Lefebvre E: Fast unfolding of communities in large networks. J. Stat. Mech. Theory Exp. 2008, P10008.(10), P10008 (12pp). doi: 10.1088/1742-5468/2008/10/P10008.

27. Lancichinetti A, Fortunato S: Community detection algorithms: A comparative analysis. Phys Rev E 2009, 80:056117.

28. Mello JF, Buzas MA: An application of cluster analysis as a method of determining biofacies. J Paleontol 1968, 42: 747–758.

29. Csárdi G, Nepusz T: The igraph software package for complex network research, InterJournal, Complex Systems 2006, 1695. http://igraph.org

